# ERGA-BGE genome of *Pinctada radiata* (Leach, 1814): one of the first Lessepsian migrants

**DOI:** 10.1101/2025.01.17.632165

**Authors:** Katerina Vasileiadou, Tereza Manousaki, Thanos Dailianis, Grigorios Skouradakis, Emmanouela Vernadou, Danae Karakasi, Astrid Böhne, Rita Monteiro, Rosa Fernández, Nuria Escudero, Genoscope Sequencing Team, Alice Moussy, Corinne Cruaud, Karine Labadie, Lola Demirdjian, Benjamin Istace, Arnaud Couloux, Patrick Wincker, Pedro H. Oliveira, Jean-Marc Aury, Francesca Floriana Tricomi, Leanne Haggerty, Fergal Martin, Chiara Bortoluzzi

## Abstract

*Pinctada radiata*, commonly known as the Gulf pearl oyster, is a species of pearl oyster found primarily in the warm waters of the Red Sea, the Persian Gulf, and parts of the Indian Ocean. *Pinctada radiata* contributes to marine ecosystems by filtering water, which helps maintain water quality and supports other marine life. This species is the first bivalve Lessepsian migrant, having migrated from the Red Sea to the Mediterranean Sea via the Suez Canal. The reference genome of *Pinctada radiata* could help identify genes enabling adaptation to varying temperatures and salinities, facilitating survival in diverse and newly colonized habitats allowing comparisons with other bivalves to uncover shared and unique genetic adaptations. Additionally, the genome could support targeted management practices and conservation initiatives, such as habitat restoration and selective breeding, ensuring the long-term sustainability of *P. radiata*. The entirety of the genome sequence was assembled into 14 contiguous chromosomal pseudomolecules. This chromosome-level assembly encompasses 0.93 Gb, composed of 220 contigs and 44 scaffolds, with contig and scaffold N50 values of 8.1 Mb and 63.8 Mb, respectively.

## Introduction

*Pinctada radiata*, or Μαργαριτοφόρο στρείδι in Greek, belongs to the *Pteriidae* family, is a marine bivalve, and is native to the Indian Ocean, Persian Gulf, and Red Sea (Aguilo-Arce et al., 2023). The taxonomic identification between the species of the family is based on the arrangement of the hinge teeth and morphological and anatomical character variation (Al-Saadi, 2013). *Pinctada radiata* was the first bivalve species recorded as a Lessepsian migrant in 1874 (Gofas et al., 2003) and is usually found in coastal habitats. The species is a protandrous hermaphrodite with developmental stages including a planktonic larval stage (Al Saadi, 2013) with good dispersal ability. There is a high economic significance to the species since it is widely cultivated for pearls, and recently it has also been farmed for food (Pafras et al., 2024). *Pinctada radiata* is not included on the IUCN Red List nor the CITES list of protected species.

*Pinctada radiata*, like most bivalves, is a filter-feeder. It has been proven to be an effective ecological indicator, as it can accumulate substantial quantities of pollutants in its tissues (Al-Madfa et al., 1998). The species can filter up to 1,500 liters of seawater and accumulate 100 times higher concentrations of pollutants than the adjacent water, therefore it is widely used in marine pollution studies, as a bio-indicator for polluted marine environments.

Developing a high-quality reference genome for *Pinctada radiata* is essential for understanding its unique genetic makeup and adaptive traits, supporting marine conservation efforts by providing insights into its resilience to polluted environments, and enhancing economic and population genomic applications through advancements in aquaculture practices and studies on its establishment as a Lessepsian migrant.

The generation of this reference resource was coordinated by the European Reference Genome Atlas (ERGA) initiative’s Biodiversity Genomics Europe (BGE) project, supporting ERGA’s aims of promoting transnational cooperation to promote advances in the application of genomics technologies to protect and restore biodiversity (Mazzoni et al., 2023).

## Materials & Methods

ERGA’s sequencing strategy includes Oxford Nanopore Technology (ONT) and/or Pacific Biosciences (PacBio) for long-read sequencing, along with Hi-C sequencing for chromosomal architecture, Illumina Paired-End (PE) for polishing (i.e. recommended for ONT-only assemblies), and RNA sequencing for transcriptome profiling, to facilitate genome assembly and annotation.

### Sample and Sampling Information

Thanos Dailianis sampled one specimen of *Pinctada radiata*, which was determined based on Manousis (2021). The specimen was identified by Katerina Vasileiadou, from the sample collected in Elounda Bay, Lasithi, Crete (Greece) on 31st May 2023. Sampling was performed under permission ΥΠΕΝ\ΔΔΔ\34284\1131 from the Ministry for Environment and Energy Secretariat General for Natural Environment & Water Directorate General for Forests & Forest Environment Directorate for Forest Management. The specimen was hand-picked. Tissues were removed from a living specimen and were flash frozen in liquid nitrogen and preserved at -80 °C until DNA extraction.

### Vouchering information

Physical reference materials for the sequenced specimen were deposited in the National History Museum of Crete https://www.nhmc.uoc.gr/ under the accession number NHMC.52.28.

Frozen reference tissue material of muscle is available from the same individual at the Biobank of the National History Museum of Crete https://www.nhmc.uoc.gr/ under the voucher ID NHMC.52.28.

An electronic voucher image of the sequenced individual is available from ERGA’s EBI BioImageArchive dataset https://www.ebi.ac.uk/biostudies/bioimages/studies/S-BIAD1012?query=ERGA under accession ID https://ftp.ebi.ac.uk/biostudies/fire/S-BIAD/012/S-BIAD1012/Files/ERGA/SAMEA114349619_1.jpg.

### Data Availability

*Pinctada radiata* and the related genomic study were assigned to Tree of Life ID (ToLID) ‘xbPinRadi1’ and all sample, sequence, and assembly information are available under the umbrella BioProject PRJEB77217. The sample information is available at the following BioSample accessions: SAMEA114349628 and SAMEA114349629.

The genome assembly is accessible from ENA under accession number GCA_964261305.1 and the annotated genome is available through the Ensembl Beta website (https://projects.ensembl.org/erga-bge/).

Sequencing data produced as part of this project are available from ENA at the following accessions: ERX12733444, ERX12733465, ERX12737196, and ERX12737197. Documentation related to the genome assembly and curation can be found in the ERGA Assembly Report (EAR) document available at https://github.com/ERGA-consortium/EARs/tree/main/Assembly_Reports/Pinctada_radiata/xbPinRadi1. Further details and data about the project are hosted on the ERGA portal at https://www.ebi.ac.uk/biodiversity/data_portal/112209.

### Genetic Information

The estimated genome size, based on ancestral taxa, is 1.15 Gb. This is a diploid genome with a haploid number of 14 chromosomes (2n=28) and unknown sex chromosomes. All information for this species was retrieved from Genomes on a Tree (Challis et al., 2023).

### DNA/RNA processing

DNA was extracted from 600 mg of muscle tissue using a conventional CTAB extraction followed by purification using Qiagen Genomic tips (Qiagen, MD, USA). A detailed protocol is available on protocols.io (https://www.protocols.io/view/hmw-dna-extraction-for-long-read-sequencing-using-bp2l694yzlqe/v1). DNA fragment size selection was performed using the Short Read Eliminator kit (PacBio). Quantification was performed using a Qubit dsDNA HS Assay kit (Thermo Fisher Scientific) and integrity was assessed in a FemtoPulse system (Agilent). DNA was stored at 4 °C until usage.

RNA was extracted from muscle (50 mg) using the RNeasy Plus Universal kit (Qiagen) following manufacturer instructions. Residual genomic DNA was removed with 6U of TURBO DNase (2 U/μL) (Thermo Fisher Scientific). Quantification was performed using a Qubit RNA HS Assay kit and integrity was assessed in a Bioanalyzer system (Agilent). RNA was stored at -80 °C.

### Library Preparation and Sequencing

Long-read DNA libraries were prepared with the SMRTbell prep kit 3.0 following manufacturers’ instructions and sequenced on a Revio system (PacBio). Hi-C libraries were generated from muscle tissue using the Arima High Coverage HiC kit (following the Animal Tissues low input protocol v01) and sequenced on a NovaSeq 6000 instrument (Illumina) with 2x150 bp read length. Poly(A) RNA-Seq libraries were constructed using the Illumina Stranded mRNA Prep, Ligation kit (Illumina) and sequenced on a NovaSeq 6000 instrument.

In total, 42x PacBio HiFi and 54x HiC data were sequenced to generate the assembly.

### Genome Assembly Methods

The genome of *Pinctada radiata* was assembled using the Genoscope GALOP pipeline (https://workflowhub.eu/workflows/1200).

Briefly, raw PacBio HiFi reads were assembled using Hifiasm v0.19.5-r593 (Cheng et al., 2021). Retained haplotigs were removed using purge_dups v1.2.5 (Guan et al., 2020) with default parameters and the proposed cutoffs. The purged assembly was scaffolded using YaHS v1.2 (Zhou et al., 2023) and assembled scaffolds were then curated through manual inspection using PretextView v0.2.5 to remove false joins and incorporate sequences not automatically scaffolded into their respective locations within the chromosomal pseudomolecules. The mitochondrial genome was assembled as a single circular contig using Oatk v1.0 (Zhou et al., 2024)and included in the released assembly.

### Genome Annotation Methods

A gene set was generated using the Ensembl Gene Annotation system (Aken et al., 2016), primarily by aligning publicly available short-read RNA-seq data from BioSamples: SAMEA114349638, SAMD00178211, SAMD00178207, SAMD00178208, SAMD00178209, SAMD00178210, SAMEA114349638 to the genome. Gaps in the annotation were filled via protein-to-genome alignments of a select set of clade-specific proteins from UniProt (Consortium, 2019), which had experimental evidence at the protein or transcript level. At each locus, data were aggregated and consolidated, prioritising models derived from RNA-seq data, resulting in a final set of gene models and associated non-redundant transcript sets. To distinguish true isoforms from fragments, the likelihood of each open reading frame (ORF) was evaluated against known metazoan proteins. Low-quality transcript models, such as those showing evidence of fragmented ORFs, were removed. In cases where RNA-seq data were fragmented or absent, homology data were prioritised, favouring longer transcripts with strong intron support from short-read data. The resulting gene models were classified into two categories: protein-coding, and long non-coding. Models that did not overlap protein-coding genes, and were constructed from transcriptomic data were considered potential lncRNAs. Potential lncRNAs were further filtered to remove single-exon loci due to their unreliability. Putative miRNAs were predicted by performing a BLAST search of miRBase (Kozomara et al., 2019) against the genome, followed by RNAfold analysis (Gruber et al., 2008). Other small non-coding loci were identified by scanning the genome with Rfam (Kalvari et al., 2018) and passing the results through Infernal (Nawrocki & Eddy, 2013). Summary analysis of the released annotation was carried out using the ERGA-BGE Genome Report ANNOT Galaxy workflow (https://workflowhub.eu/workflows/1096).

## Results

### Genome Assembly

The genome assembly has a total length of 931,129,676 bp in 44 scaffolds including the mitogenome (Figures 1 & 2), with a GC content of 35.4%. The assembly has a contig N50 of 8,065,000 bp and L50 of 34 bp and a scaffold N50 of 63,836,525 bp and L50 of 6 bp. The assembly has a total of 176 gaps, totaling 24.2 kb in cumulative size. The single-copy gene content analysis using the Eukaryota database with BUSCO (Manni et al., 2021) resulted in 98.1% completeness (97.3% single and 0.8% duplicated). 67.7% of reads k-mers were present in the assembly and the assembly has a base accuracy Quality Value (QV) of 61.7 as calculated by Merqury (Rhie et al., 2020).

**Figure 1.**
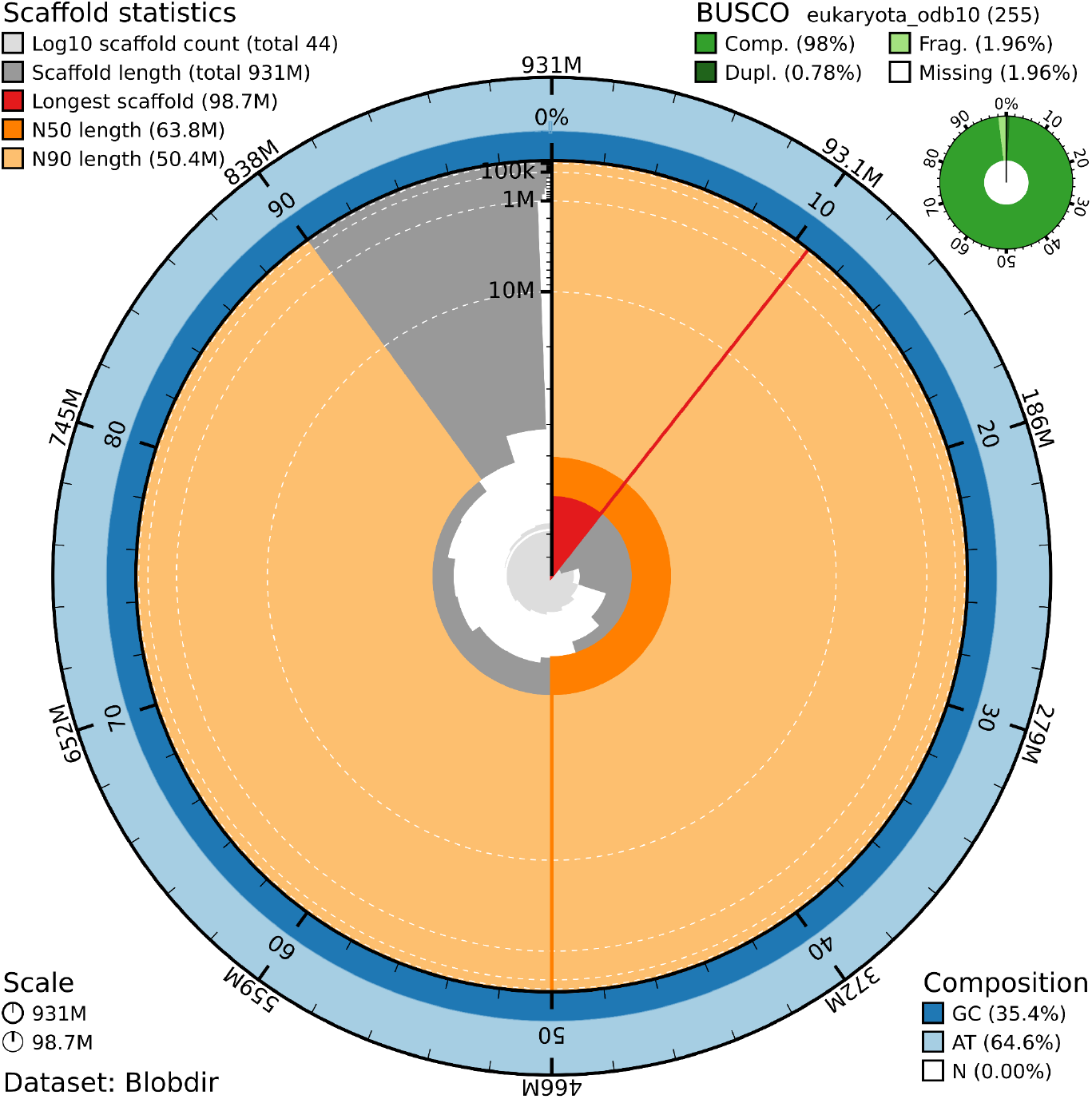
Snail plot summary of assembly statistics. The main plot is divided into 1,000 size-ordered bins around the circumference, with each bin representing 0.1% of the 931,129,676 bp assembly including the mitochondrial genome. The distribution of sequence lengths is shown in dark grey, with the plot radius scaled to the longest sequence present in the assembly (98.7 Mb, shown in red). Orange and pale-orange arcs show the scaffold N50 and N90 sequence lengths (63,836,525 and 50,368,007 bp), respectively. The pale grey spiral shows the cumulative sequence count on a log-scale, with white scale lines showing successive orders of magnitude. The blue and pale-blue area around the outside of the plot shows the distribution of GC, AT, and N percentages in the same bins as the inner plot. A summary of complete, fragmented, duplicated, and missing BUSCO genes found in the assembled genome from the Eukaryota database (odb10) is shown in the top right.

**Figure 2.**
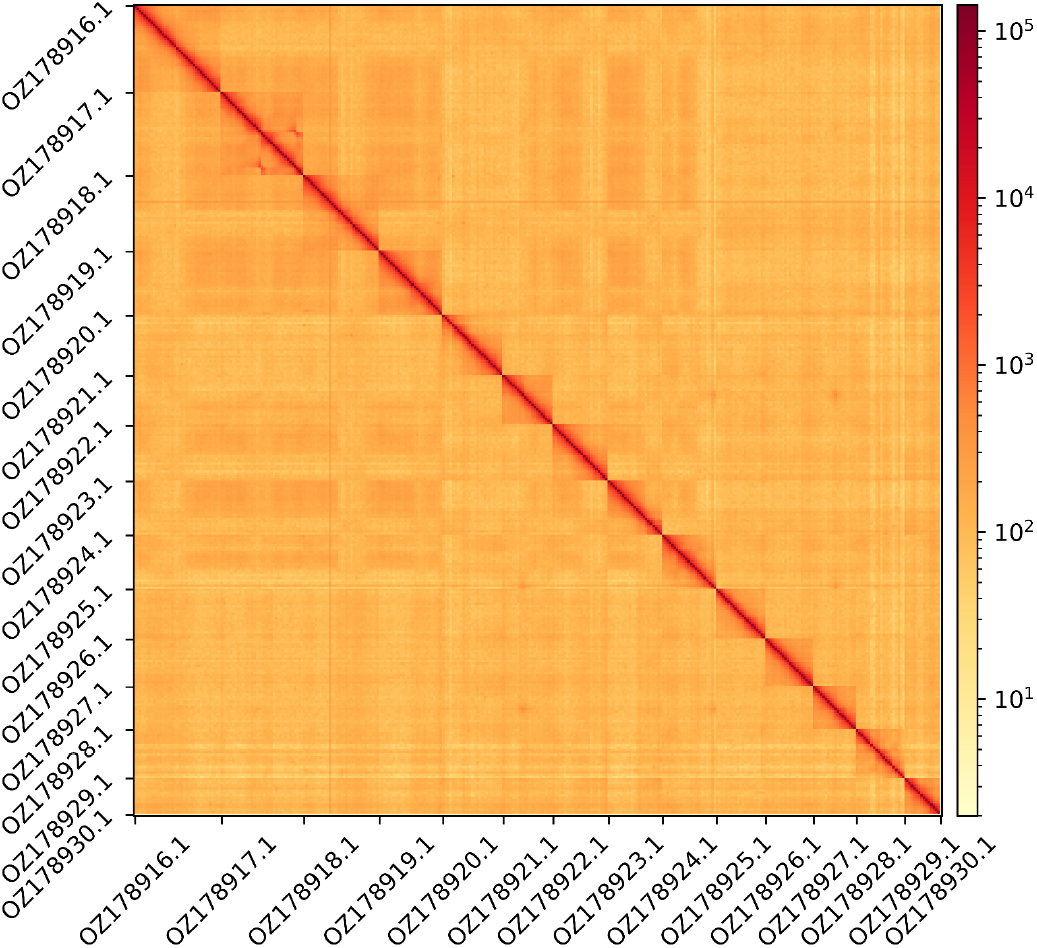
Hi-C contact map showing spatial interactions between regions of the genome. The diagonal corresponds to intra-chromosomal contacts, depicting chromosome boundaries. The frequency of contacts is shown on a logarithmic heatmap scale. Hi-C matrix bins were merged into a 25 kb bin size for plotting. Due to space constraints on the axes, only the GenBank names of the 14th largest chromosomes and the mitochondrial genome (GenBank name: OZ178930.1) are shown.

### Genome Annotation

The genome annotation consists of 23,106 protein-coding genes with associated 40,414 transcripts, in addition to 17,566 non-coding genes (Table 1). Using the longest isoform per transcript, the single-copy gene content analysis using the Eukaryota database with BUSCO resulted in 98.1% completeness. Using the OMAmer Metazoa-v2.0.0.h5 database for OMArk (Nevers et al., 2025) resulted in 96.2% completeness and 57.9% consistency (Table 2).

**Table 1.** Statistics from assembled gene models.

|  | No. genes | No. transcripts | Mean gene length (bp) | No. single-exon genes | Mean exons per transcript |
| --- | --- | --- | --- | --- | --- |
| mRNA | 23,106 | 40,414 | 16,303 | 776 | 8.9 |
| pseudogene | 0.0 | 0.0 | 0.0 | 0.0 | 0.0 |
| snoRNA | 33 | 33 | 117 | 33 | 1.0 |
| lncRNA | 6,862 | 8,699 | 6,773 | 4 | 2.4 |
| miRNA | 0.0 | 0.0 | 0.0 | 0.0 | 0.0 |
| snRNA | 226 | 226 | 147 | 226 | 1.0 |
| rRNA | 159 | 159 | 254 | 159 | 1.0 |
| scRNA | 1 | 1 | 127 | 1 | 1.0 |
| Other ncRNA | 10,285 | 10,285 | 63-72 | 10,285 | 1.0 |

**Table 2.**
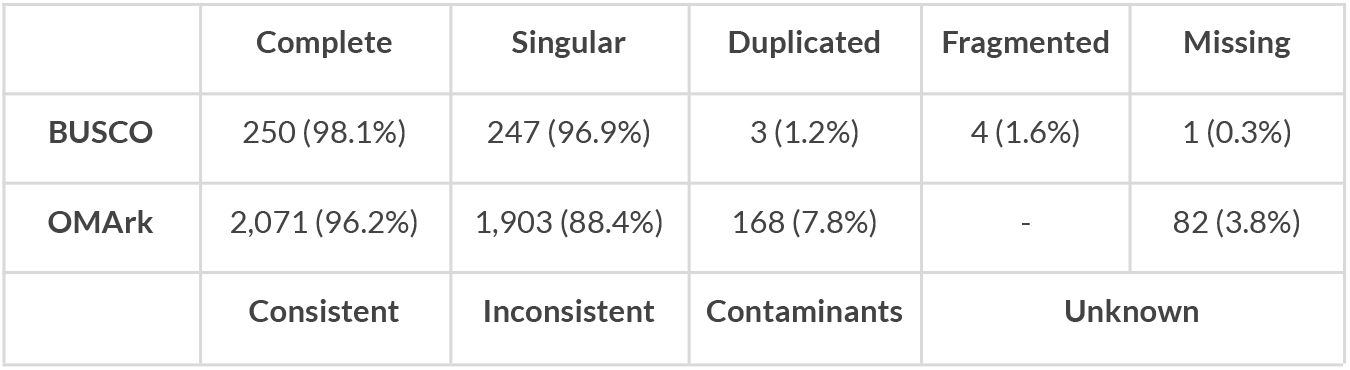
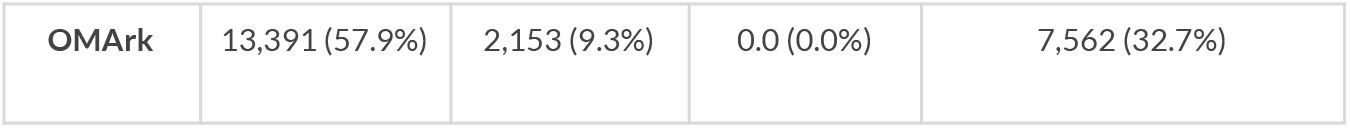
Annotation completeness and consistency scores calculated by BUSCO run in protein mode (eukaryota_odb10) and OMArk (Metazoa-v2.0.0.h5)

|  | Complete | Singular | Duplicated | Fragmented | Missing |
| --- | --- | --- | --- | --- | --- |
| BUSCO | 250 (98.1%) | 247 (96.9%) | 3 (1.2%) | 4 (1.6%) | 1 (0.3%) |
| OMArk | 2,071 (96.2%) | 1,903 (88.4%) | 168 (7.8%) | - | 82 (3.8%) |
|  | Consistent | Inconsistent | Contaminants | Unknown |  |

|  |  |  |  |  |
| --- | --- | --- | --- | --- |
| OMark | 13,391 (57.9%) | 2,153 (9.3%) | 0.0 (0.0%) | 7,562 (32.7%) |

## Acknowledgements

We would like to acknowledge the assembly reviewer, Michael Paulini, from the Wellcome Sanger Institute, United Kingdom.

## Conflict of Interest

The authors declare no conflict of interest related to this study. The funding sources had no involvement in the study design, collection, analysis, or interpretation of data; in the writing of the manuscript; or in the decision to submit the article for publication. All authors have participated sufficiently in the work to take public responsibility for the content and agree to the submission of this manuscript.

## Funder Information

This work received funding from Biodiversity Genomics Europe (Grant no.101059492), which is funded by Horizon Europe under the Biodiversity, Circular Economy and Environment call (REA.B.3); co-funded by the Swiss State Secretariat for Education, Research and Innovation (SERI) under contract numbers 22.00173 and 24.00054; and by the UK Research and Innovation (UKRI) under the Department for Business, Energy and Industrial Strategy’s Horizon Europe Guarantee Scheme. This work was supported by the Genoscope, the Commissariat à l’Energie Atomique et aux Énergies Alternatives (CEA), France Génomique (ANR-10-INBS-09-08) and the exploratory research programme “ATLASea: Atlas of marine genomes” and its targeted project SEQ-Sea (ANR-22-EXAT-0003-SEQ-Sea).

## Author Contributions

KV and TM coordinated the project; TD, GS, and EV collected the species; TD, GS, and EV identified the species; KV, TM, and DK sampled and preserved biological material and provided metadata; RM, NE, RF, and AsB provided support in sampling, shipping of biological material, metadata collection, and management; the GST extracted DNA, prepared libraries, and performed sequencing under the supervision of AM, CC, KL, PHO and PW; LD, BI, AC and JMA performed genome assembly and curation; LH, SS, and FM performed genome annotation; CB generated the analysis and report. All authors contributed to the writing, review, and editing of this genome note and read and approved the final version. This work is part of the species assigned to Genoscope, which was instrumental in the wet lab, sequencing, and assembly processes, and represents a key contribution to BGE’s outputs.

## Author Information

Members of the Genoscope Sequencing Team are listed here: https://zenodo.org/records/1461149.

## Notes

https://www.ebi.ac.uk/ena/browser/view/PRJEB77217

### Competing Interest Statement

The authors have declared no competing interest.

### Summary of Updates

This version of the manuscript has been revised to update the Genome Annotation Methods.

## Literature Cited

Aguilo-Arce, J., Ferragut, J., Png-González, L., Carbonell, A., & Capa, M. (2023). First genetic survey on the invasive rayed pearl oyster Pinctada radiata (Leach, 1814) populations of the Balearic Islands (Western Mediterranean). Mediterranean Marine Science, 24(3), 666–678.

Aken, B. L., Ayling, S., Barrell, D., Clarke, L., Curwen, V., Fairley, S., Fernandez Banet, J., Billis, K., García Girón, C., & Hourlier, T. (2016). The Ensembl gene annotation system. Database, 2016, baw093.

Al Saadi, A. (2013). Population structure and patterns of genetic variation in a pearl oyster (Pinctada radiata) native to the Arabian Gulf [PhD Thesis, Queensland University of Technology]. https://eprints.qut.edu.au/62410/

Al-Madfa, H., Abdel-Moati, M. A. R., & Al-Gimaly, F. H. (1998). Pinctada radiata (Pearl Oyster): A bioindicator for metal pollution monitoring in the Qatari waters (Arabian Gulf). Bulletin of Environmental Contamination and Toxicology, 60, 245–251.

Challis, R., Kumar, S., Sotero-Caio, C., Brown, M., & Blaxter, M. (2023). Genomes on a Tree (GoaT): A versatile, scalable search engine for genomic and sequencing project metadata across the eukaryotic tree of life. Wellcome Open Research, 8, 24.

Cheng, H., Concepcion, G. T., Feng, X., Zhang, H., & Li, H. (2021). Haplotype-resolved de novo assembly using phased assembly graphs with hifiasm. Nature Methods, 18(2), 170–175.

Consortium, U. (2019). UniProt: A worldwide hub of protein knowledge. Nucleic Acids Research, 47(D1), D506–D515.

Gofas, S., Zenetos, A., Gibson, R. N., & Atkinson, R. J. A. (2003). Exotic molluscs in the Mediterranean basin: Current status and perspectives. Oceanography and Marine Biology: An Annual Review, 41, 237–277.

Gruber, A. R., Lorenz, R., Bernhart, S. H., Neuböck, R., & Hofacker, I. L. (2008). The vienna RNA websuite. Nucleic Acids Research, 36(Suppl_2), W70–W74.

Guan, D., McCarthy, S. A., Wood, J., Howe, K., Wang, Y., & Durbin, R. (2020). Identifying and removing haplotypic duplication in primary genome assemblies. Bioinformatics, 36(9), 2896–2898.

Kalvari, I., Nawrocki, E. P., Argasinska, J., Quinones-Olvera, N., Finn, R. D., Bateman, A., & Petrov, A. I. (2018). Non-Coding RNA Analysis Using the Rfam Database. Current Protocols in Bioinformatics, 62(1), e51.

Kozomara, A., Birgaoanu, M., & Griffiths-Jones, S. (2019). miRBase: From microRNA sequences to function. Nucleic Acids Research, 47(D1), D155–D162.

Manni, M., Berkeley, M. R., Seppey, M., Simão, F. A., & Zdobnov, E. M. (2021). BUSCO update: Novel and streamlined workflows along with broader and deeper phylogenetic coverage for scoring of eukaryotic, prokaryotic, and viral genomes. Molecular Biology and Evolution, 38(10), 4647–4654.

Manousis, T. (2021). Hellenic Conches. Harxheim: Conchbooks, 609 pp. ISBN 978-3-948603-17-5.

Nawrocki, E. P., & Eddy, S. R. (2013). Infernal 1.1: 100-fold faster RNA homology searches. Bioinformatics, 29(22), 2933–2935.

Nevers, Y., Warwick Vesztrocy, A., Rossier, V., Train, C.-M., Altenhoff, A., Dessimoz, C., & Glover, N. M. (2025). Quality assessment of gene repertoire annotations with OMArk. Nature Biotechnology, 43(1), 124–133.

Pafras, D., Theocharis, A., Kondylatos, G., Conides, A., & Klaoudatos, D. (2024). Population Biology of the Non-Indigenous Rayed Pearl Oyster (Pinctada radiata) in the South Evoikos Gulf, Greece. Diversity, 16(8), 460.

Rhie, A., Walenz, B. P., Koren, S., & Phillippy, A. M. (2020). Merqury: Reference-free quality, completeness, and phasing assessment for genome assemblies. Genome Biology, 21(1), 245.

Zhou, C., Brown, M., Blaxter, M., Consortium, D. T. of L. P., McCarthy, S. A., & Durbin, R. (2024). Oatk: A de novo assembly tool for complex plant organelle genomes. bioRxiv, 2024–10.

Zhou, C., McCarthy, S. A., & Durbin, R. (2023). YaHS: Yet another Hi-C scaffolding tool. Bioinformatics, 39(1), btac808.

